# The *mir-35* family links maternal germline sex to embryonic viability in *C. elegans*

**DOI:** 10.1101/516492

**Authors:** Lars Benner, Katherine Prothro, Katherine McJunkin

**Affiliations:** NIDDK Intramural Research Program, National Institutes of Health, Bethesda, MD 20892

**Keywords:** sex determination, microRNAs, embryogenesis, XO lethality, germline sex determination

## Abstract

The germline sex determination pathway in *C. elegans* determines whether germ cells develop as oocytes or sperm, with no previously known effect on viability. The *mir-35* family of microRNAs are expressed in the *C. elegans* germline and embryo and are essential for both viability and normal hermaphroditic sex determination, preventing aberrant male gene expression in XX hermaphrodite embryos. Here we show that combining feminizing mutations with partial loss of function of the *mir-35* family results in enhanced penetrance embryonic lethality that preferentially kills XO animals. This lethal phenotype is due to altered signaling through the germline sex determination pathway, and maternal germline feminization is sufficient to induce enhanced lethality. These findings reveal a surprising pleiotropy of sperm-fate promoting pathways on organismal viability. Overall, our results demonstrate an unexpectedly strong link between sex determination and embryonic viability, and suggest that in wild type animals, *mir-35* family members buffer against misregulation of pathways outside the sex determination program, allowing for clean sex reversal rather than deleterious effects of perturbing sex determination genes.

## Introduction

MicroRNAs are a class of endogenous 22-23-nucleotide RNAs that repress expression of complementary target mRNAs to govern diverse processes in essentially all complex eukaryotes. The seed region (nucleotides 2-7) of a microRNA is the most important portion of the sequence for determining target specificity (Bartel 2009). MicroRNAs which share the same seed sequence are classified as a “family” because they can potentially bind and redundantly regulate the same set of target mRNAs.

The *mir-35-42* microRNA family is abundantly expressed in oocytes and early embryos and is essential for *C. elegans* embryonic development (Alvarez-Saavedra and Horvitz 2010; Wu *et al.* 2010). The *mir-35* family consists of eight members (*mir-35-42*) which reside in two loci (*mir-35-41* and *mir-42-44*). While deletion of all eight *mir-35-42* microRNA genes results in completely penetrant embryonic or early larval lethality (Alvarez-Saavedra and Horvitz 2010), strains which carry a deletion affecting only the *mir-35-41* cluster display a weaker temperature-sensitive lethal phenotype with low penetrance of lethality at 15°C or 20°C and nearly complete lethality at 25°C (Alvarez-Saavedra and Horvitz 2010).

Since *mir-35-41* deletion mutants can bypass embryonic lethality at permissive temperature, this genetic setting is tractable for understanding which pathways are deregulated in *mir-35* family loss of function (Liu *et al.* 2011; Massirer *et al.* 2012; McJunkin and Ambros 2014). We previously demonstrated that the most strongly upregulated genes in hermaphrodite *mir-35-41(nDf50)* embryos are targets of the sex determination pathway that are normally repressed in hermaphrodites and highly expressed in males (McJunkin and Ambros 2017). We found that this aberrant upregulation of male-specific gene expression was driven by derepression of two *mir-35-41* target genes encoding RNA binding proteins. One of these genes*, <u>SUP</u>pressor-26* (*sup-26*), had previously been identified as a male-promoting modulator of the sex determination pathway (Manser *et al.* 2002; Mapes *et al.* 2010) while the other gene, *<u>NHL</u> domain containing-2* (*nhl-2*), was not previously implicated in sex determination.

The fact that *mir-35-41* regulate sex determination is surprising since they are maternally contributed (as well as zygotically expressed) and act partially by maternal effect. Because sex must be determined zygotically, we postulate that the *mir-35* family prevents premature sex-specific gene expression. This role of the *mir-35* family as a developmental timer is analogous to the roles of other essential microRNA families such as *lin-4*, *let-7* and the *let-7* sisters, each of which is temporally expressed to drive forward developmental transitions during larval development (Feinbaum and Ambros 1999; Reinhart *et al.* 2000; Abbott *et al.* 2005).

Sex determination is a rapidly evolving process that is governed by a diversity of genetic mechanisms across animal phylogeny. Because of its evolutionary plasticity, sex determination is well suited for regulation by microRNAs, whose target-specificity is defined by a minute genomic space (6-7 nucleotides), and thus is rapidly evolving.

In the *C. elegans* hermaphrodite, a female soma harbors a germline which produces sperm during larval development and oocytes throughout adulthood. The female somatic developmental program is initiated in animals with two X chromosomes (XX) by the dosage compensation complex (DCC), which acts to silence transcription of X-linked genes by two-fold. The DCC also silences the master regulator of sex determination, <u>her</u>maphroditization of XO animals-1 (*her-1*), by forty-fold (Meyer 2005) (Figure 1A). *Her-1* encodes a diffusible ligand that binds to and inhibits the Patched-like transmembrane receptor <u>Tra</u>nsformer-2 (TRA-2) (Wolff and Zarkower 2008). Thus, low levels of HER-1 in hermaphrodites allow proteolytic activation of TRA-2. Active TRA-2 inhibits the intracellular complex of FEM proteins (<u>FEM</u>inization of XX and XO animals), whose activity antagonizes the transcription factor <u>Tra</u>nsformer-1 (TRA-1). Thus, TRA-1 is licensed to repress its multiple target genes, ensuring proper female somatic development. Conversely, in males, only one X chromosome is present (XO), and dosage compensation is off. This permits high levels of HER-1 expression, which leads to an active FEM complex, and high levels of TRA-1 target genes, which carry out the male-specific developmental program. We previously found that the targets of *mir-35-41* act at multiple levels in this pathway – both upstream and downstream of *her-1* – to drive aberrant male-like gene expression in *mir-35-41(nDf50)* mutant hermaphrodites (McJunkin and Ambros 2017).

**Figure 1.**
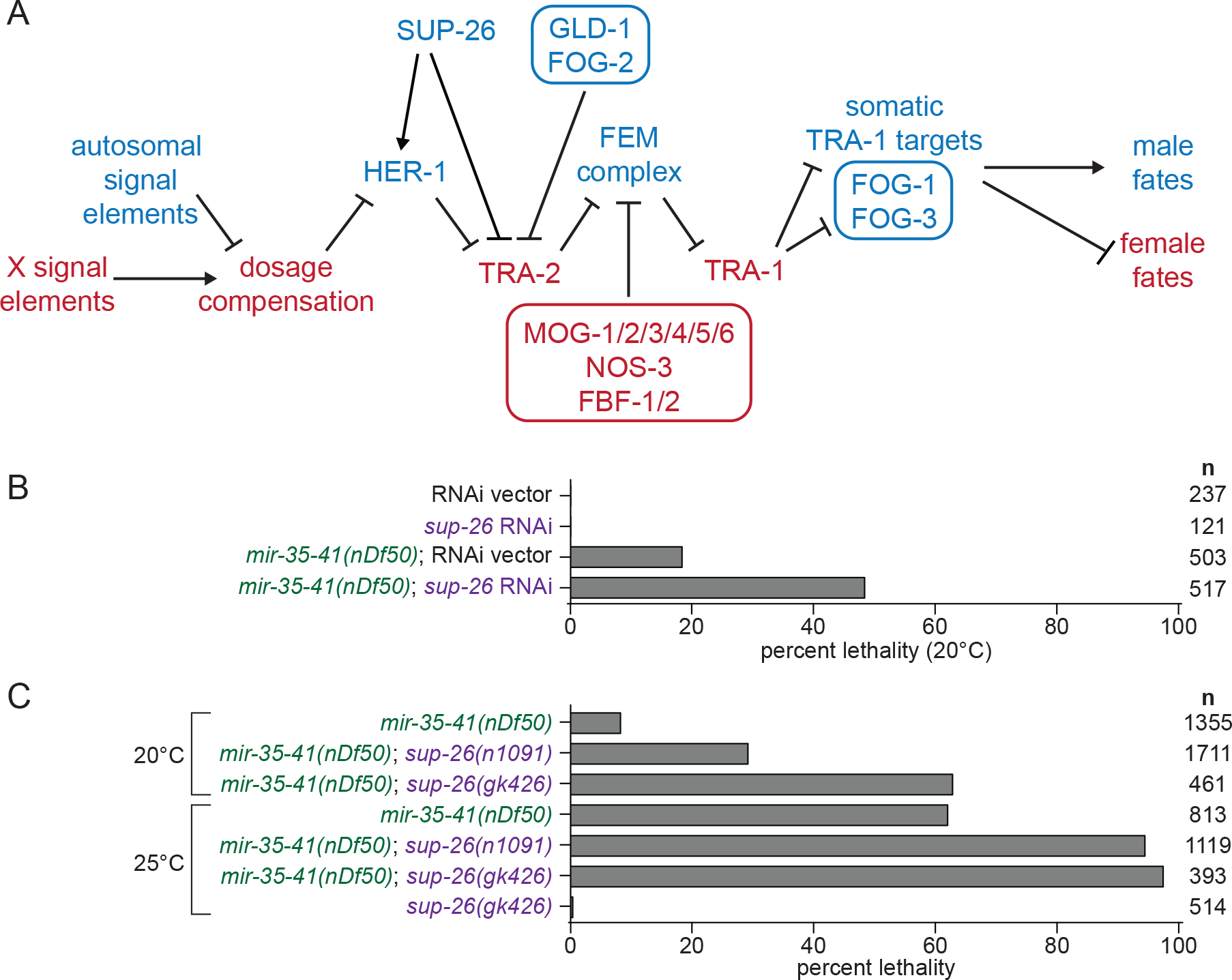
SUP-26 promotes embryonic viability in *mir-35-41(nDf50)*. A) A schematic of the principal genetic interactions of sex determination pathway genes in *C. elegans*. Genes in blue are highly active in the male soma and germline, and the spermatogenic germ cells in hermaphrodites. Genes in red are highly active specifically in the hermaphrodite soma and oogenic germ cells. Germline-specific factors are in boxes. B) Percent lethality at embryonic or early larval stages. *Sup-26(RNAi)* has no effect on wild type, but induces synthetic lethality in *mir-35-41(nDf50)*. C) Percent lethality at embryonic or early larval stages. *Sup-26(lf)* alleles enhance lethality in *mir-35-41(nDf50)*.

Sex determination of the germ cell lineage is governed by the same core pathway as the soma, but additional players act to transduce the upstream signal to produce a sperm or oocyte cell fate. Additionally, in hermaphrodites, temporal regulation of the sex determination pathway in the germline allows for the production of sperm during larval development followed by production of oocytes in the germline of adults. In both males and hermaphrodites, TRA-1 controls germ cell fate by regulating <u>F</u>eminization <u>o</u>f <u>G</u>ermline-1 and −3 (*fog-1* and *fog-3*) (Ellis and Schedl 2007) (Figure 1A). When expressed, both FOG-1 (a cytoplasmic polyadenylation element binding protein) and FOG-3 (a Tob family protein) promote male cell fate (spermatogenesis). In hermaphrodites, additional players act to transiently repress *tra-2* translation early in development to promote spermatogenesis, while other factors repress *fem-3* translation later to promote the switch to oogenesis. These factors include the *tra-2* repressor *defective in <u>G</u>erm<u>L</u>ine <u>D</u>evelopment 1* (GLD-1) with its partner FOG-2 (Figure 1A). Negative regulators of *fem-3* translation include *<u>N</u>anOS-<u>r</u>elated 3* (*nos-3*), *<u>F</u>em-3 mRNA <u>B</u>inding <u>F</u>actor-1* and −*2* (*fbf-1* and *fbf-2*) and *<u>M</u>asculinization <u>O</u>f <u>G</u>ermline-1-6* (*mog-1* through *mog-6*).

The genes encoding the DCC are essential in XX animals because of their function in dosage compensation of X-linked genes; conversely, overexpression of the DCC in XO animals is lethal. In contrast, sex determination genes downstream of dosage compensation affect sexual dimorphism but are not essential. Generally, their mutation causes sex reversal, but not lethality. For instance, null alleles of *tra-2* or *tra-1* result in XX animals that undergo male development, while null alleles of *her-1* or *fem* genes cause somatic feminization of XO animals. Mutations in germline-specific sex determination genes can result in germline-specific sex reversal, and are not known to affect viability.

In this work, we examine how the sex determination pathway genetically interacts with *mir-35-41* mutant lethality phenotypes. We find that *mir-35-41* prevents lethality in feminized backgrounds, surprisingly demonstrating a role for male-promoting genes (e.g. *her-1*) in hermaphrodite viability. Furthermore, we find that the synthetic lethality of *mir-35-41* deletion with feminizing mutations preferentially affects XO animals. Finally, we observe that these synthetic lethal effects are mediated by the germline-specific module of the sex determination pathway via a maternal effect. To our knowledge this is the first description of the germline sex determination pathway modulating viability.

## Materials and Methods

### *C. elegans* culture and RNAi

*C. elegans* were maintained on NGM seeded with HB101. Strains were kept at 15°C or 20°C for 72 hours or 25°C for 48 hours prior to beginning experiments conducted at the respective temperature. For quantification of lethality, single hermaphrodites were placed on individual 3cm NGM plates for approximately 24 hours. The single hermaphrodites were moved to a fresh plate each day. Approximately 24 hours after removal of the parent, larvae were counted and scored as live or dead. For *him-8* strains and crosses where males and hermaphrodites were quantified, dead L1 larvae and dead embryos were counted 24 hours after removal of the parent; surviving progeny were scored as male and hermaphrodite the following day, 48 hours after removal of the parent.

For RNAi experiments, plates were supplemented with 1ug/ml IPTG, and were poured no more than two weeks prior to use, and stored at 4°C. Gravid adults were placed on RNAi food either for 48 hours (at 25°C) or for 72 hours (at 20°C) before single L4 progeny were chosen and placed individually on RNAi plates for progeny quantification as described above. For *tra-1* and *tra-2* RNAi, a *lag-2::GFP* reporter (*qIs56*) was used to select L3 hermaphrodites. These L3s were placed on RNAi at 20°C for 24h; after 24h, these animals were young adults and were shifted to 25°C, and their progeny were scored for lethality as above. The RNAi plasmids targeting *sup-26, tra-1* and *tra-2* are from the Vidal ORFeome RNAi library (Rual *et al.* 2004).

Alleles used were *mir-35-41(nDf50), sup-26(n1091), sup-26(gk426), him-8(e1489), lon-2(e678), tra-2(e2020gf), tra-2(e1095null), her-1(hv1y101null), xol-1(y9null), fem-3(e1996null)*, *fog-2(q71null)*, *fog-1(q325null)*, and *fog-3(q470null)*. The balancer qC1 marked with *qIs26 (rol-6(su1006), lag-2::GFP)* was used as a balancer for *sup-26(lf)* alleles in the *mir-35-41(nDf50)* background. The *mIn1* balancer marked with *mIs14* (*myo2::GFP, pes-10::GFP, F2B7.9::GFP)* was used to balance *mir-35-41(nDf50)* and *tra-2* alleles; *mIn1* provided *mir-35-41* expression in experiments where the maternal or zygotic role of *mir-35-41* was assessed*. Fem-3(e1996null)* was balanced by *nT1* with *qIs51* (*myo2::GFP, pes-10::GFP, F2B7.9::GFP)*. The X-linked mCherry transgene used to score karyotype in *fem-3(null)* crosses was oxTi421 [*eft-3p::mCherry::tbb-2* 3’UTR + *Cbr-unc-119*(+)].

For *lon-2* crosses, one L4 hermaphrodite was mated with five males. Progeny were counted each day as described above. However, to minimize the self-progeny included in quantification, all progeny counted prior to the appearance of the first male progeny were excluded. In addition, all progeny counted on the same day as the first male progeny were excluded. For example, if males were first present on day two, then only progeny from day three and later are shown. Thus, the vast majority of animals included in quantification are cross progeny, and the rare Lon hermaphrodites likely represent self-progeny.

For *mir-35-41(nDf50); fog-1(null)* and *mir-35-41(nDf50); fog-3(null)* crosses and their *mir-35-41(nDf50)* control, all parental males were homozygous for an autosomal GFP transgene (*qIs56*). In these crosses, only live progeny containing *qIs56* were counted (indicating that they are cross progeny). This eliminates the risk of counting self progeny, which is only a concern in the case of the control *mir-35-41(nDf50)* cross.

### *Tra-2* genetic experiments

*Tra-2(e2020gf)* is dominant, and prevents the development of sperm in XX animals, resulting in a male-female (rather than hermaphrodite) strain. To assess the effect of *tra-2(gf)* on *mir-35-41(nDf50)* in a pure population of XX animals, females or hermaphrodites were crossed to XX males to yield 100% XX progeny. XX males were generated by crossing *tra-2(null)* and *xol-1(null)* into to *mir-35-41(nDf50)*. *xol-1(null)* was included because *tra-2(null)* XX pseudomales are unable to mate when *xol-1* is wild type. *Mir-35-41(nDf50) tra-2(null);xol-1(null)* XX males were crossed to either a *tra-2(gf)* female or a *mir-35-41(nDf50) tra-2(gf)* female or a *mir-35-41(nDf50)* hermaphrodite. Thus, progeny are all XX, and either *mir-35-41^+/nDf50^tra-2^e2020gf/null^*;*xol-1^+lnull^* or *mir-35-41^nDf50/nDf50^tra-2^e2020gf/null^;xol-1^+/null^* or *mir-35-41^nDf50/nDf50^tra-2^+/null^;xol-1^+/null^*. Since *mir-35-41(nDf50), tra-2(null) and xol-1(null)* are all recessive, and *tra-2(e2020gf)* is dominant, the maternal genotypes highlight the functional genotypic differences. DAPI staining was performed on unmated adults in the first 24h of gravidity by fixing in 95% ethanol for five minutes, followed by mounting in Prolong Diamond antifade mountant with DAPI (ThermoFisher).

To assess the impact of *tra-2(e2020gf)* on males, a *tra-2(e2020gf)* or *mir-35-41(nDf50) tra-2(e2020gf)* female was crossed to *mir-35-41(nDf50)* males. Here, all progeny are cross-progeny because the females are self-sterile. The control, a *mir-35-41(nDf50)* hermaphrodite crossed to males of the same genotype, could potentially produce hermaphrodite self-progeny if the cross is inefficient, spuriously decreasing the apparent proportion of males. However, these data do not appear to be affected by this caveat since the proportion of males observed in the *mir-35-41(nDf50)* strain is near 50%, and is higher than in the male-female *mir-35-41(nDf50) tra-2(e2020gf)* strain (due to male lethality in the latter).

Strains are available upon request. The authors affirm that all data necessary for confirming the conclusions of the article are present within the article, figures, and tables.

## Results

### *mir-35-41* promotes viability in feminized genetic backgrounds

During the course of our previous work, we discovered that *sup-26*, a previously-characterized negative regulator of *tra-2* translation (Mapes *et al.* 2010), is required for the aberrantly masculinized phenotypes observed in *mir-35-41(nDf50)* animals and that *sup-26* acts both upstream and downstream of *her-1* in this process (Figure 1A). Surprisingly, although *sup-26(lf)* suppressed most phenotypic aspects of masculinized development in *mir-35-41(nDf50)*, knockdown or mutation of *sup-26(lf)* strongly enhanced the embryonic lethality phenotype, at both permissive and restrictive temperature (Figure 1B-C).

Our previous work showed that SUP-26 binds to hundreds of other RNA targets in addition to *tra-2*, and thus likely modulates other pathways in addition to sex determination (McJunkin and Ambros 2017). To determine whether the enhanced lethality in *mir-35-41(nDf50)*; *sup-26(lf)* was due to the effects of *sup-26* on sex determination, we examined the penetrance of *mir-35-41(nDf50)* lethality in other sex determination mutants. First we examined another feminizing mutation, *tra-2(e2020gf)*, which should partially recapitulate the effect of *sup-26(lf)* on the sex determination pathway since this *tra-2(e2020gf)* allele deletes the primary binding site of SUP-26 (Doniach 1986; Goodwin *et al.* 1993; Mapes *et al.* 2010). The sex determination phenotype of *tra-2(e2020gf)* is stronger than that of *sup-26(lf)* because GLD-1 also binds the 3’UTR element deleted in *tra-2(e2020gf)* (Doniach 1986; Jan *et al.* 1999); thus these mutants display germline feminization, requiring a cross to XX males to examine viability of XX progeny (Figure 2A) (see Methods). Like *sup-26(lf)*, *tra-2(e2020gf)* strongly enhanced *mir-35-41(nDf50)* lethality in XX animals (Figure 2A). The effect of *tra-2(e2020gf)* was further enhanced by *sup-26(RNAi)* (Figure 2A), possibly due to additional targets of SUP-26 in sex determination (McJunkin and Ambros 2017). Enhanced lethality is unlikely to be due to gross oocyte defects since germline development and mature oocytes appear normal in all maternal genotypes (Figure S1). Another feminizing mutation, deletion of *her-1*, also strongly enhanced *mir-35-41(nDf50)* lethality (Figure 2B).

**Figure 2.**
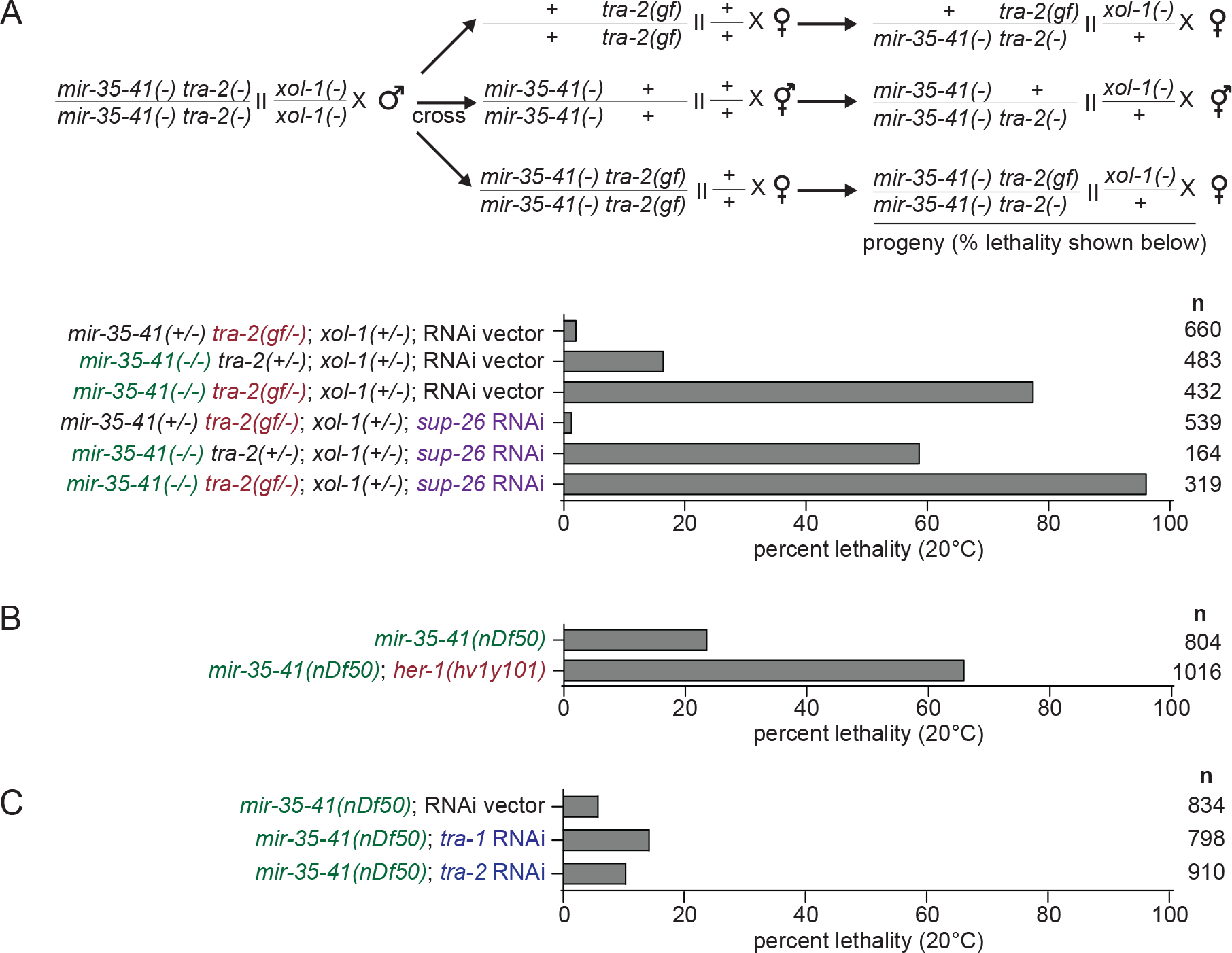
Feminizing mutations enhance lethality in *mir-35-41(nDf50)* XX animals. A) Top: schematic of crosses performed to generate progeny scored in graph. All males were XX males generated via null mutations in *tra-2* and *xol-1*. XX males were crossed to XX hermaphrodites or females of the three indicated genotypes. Bottom: percent lethality of progeny of crosses illustrated in schematic. The crosses were conducted on RNAi plates, so RNAi affects the maternal and zygotic contribution of *sup-26* in the progeny. *Sup-26(RNAi)* and *tra-2(e2020gf)* synthetically enhance XX lethality in *mir-35-41(nDf50)*. Alleles used are *tra-2(e1095lf)*, *tra-2(e2020gf)*, *xol-1(y9)*. B-C) Percent lethality in *mir-35-41(nDf50)* combined with *her-1(null)* or *tra-1* or *tra-2* RNAi.

Masculinizing treatments, such as *tra-1(RNAi)* and *tra-2(RNAi)* had a very mild effect on *mir-35-41(nDf50)* lethality (Figure 2C), and this was not due to low potency of RNAi since surviving animals displayed transformed and intersex phenotypes (Figure S2). In summary, all feminizing genetic backgrounds examined led to a strong enhancement of lethality in the *mir-35-41(nDf50)* XX animals, while masculinizing mutations did not. This phenotypic effect of feminizing mutations in XX animals is surprising since XX animals are already somatically female. Thus these results demonstrate a cryptic role for *her-1* in XX animals, where *her-1* is generally thought to be minimally expressed and biologically inactive.

### Males are preferentially affected by lethality in feminized *mir-35* family mutants

In all experiments described so far, broods of XX animals were scored for viability. We wondered whether the genetic enhancement of *mir-35-41(nDf50)* lethality by feminizing mutations was sex-specific. To analyze the impact of *mir-35-41(nDf50)* and *mir-35-41(nDf50);sup-26(lf)* lethality on males, a mutation that causes a high incidence of males (*him-8(e1489)*) through impaired X chromosome segregation was introduced to *mir-35-41(nDf50)* (Hodgkin *et al.* 1979). The *him-8* strain had 5.0% embryonic/early larval lethality, and segregated 37.5% males (among the surviving progeny) (Figure 3A). Both males and hermaphrodites were affected by increased lethality in the *mir-35-41(nDf50)*; *him-8(e1489)* background: the proportions of both populations were lower (in favor of dead embryos and larvae of unknown sex) (Figure 3A, compare first bar to third bar). Of the surviving animals in *mir-35-41(nDf50);him-8(e1489)* populations, the proportion of males (31.8%) was slightly lower than *him-8* alone, indicating that lethality due to *mir-35-41(nDf50)* impacted males slightly more than hermaphrodites (Figure 3A) (McJunkin and Ambros 2017). When *mir-35-41(nDf50)*; *him-8(e1489)* was treated with *sup-26(RNAi)*, lethality was further increased in both sexes, and even fewer males (21.6%) were present in the surviving progeny (Figure 3A). Next, we examined the *tra-2(e2020gf)* genetic background. The sex determination phenotype of *tra-2(e2020gf)* does not affect all tissues equally, primarily affecting the XX germline (Doniach 1986). As a result, *tra-2(e2020gf)* produces XX females and XO males, allowing for the sex of progeny to be scored by morphology (Figure 3B). Like *sup-26(RNAi)*, *tra-2(e2020gf)* enhanced overall lethality in *mir-35-41(nDf50)*, and a smaller proportion of males were observed among the surviving progeny (Figure 3B). As observed for XX broods, these phenotypes were further enhanced when *sup-26(RNAi)* was performed in the *tra-2(e2020gf)* background, suggesting that SUP-26 acts through other targets in addition to *tra-2* (Figure 3B). Together these results suggest that feminizing mutations enhance lethality of both sexes when *mir-35* family function is compromised, but preferentially enhance the lethality among males.

**Figure 3.**
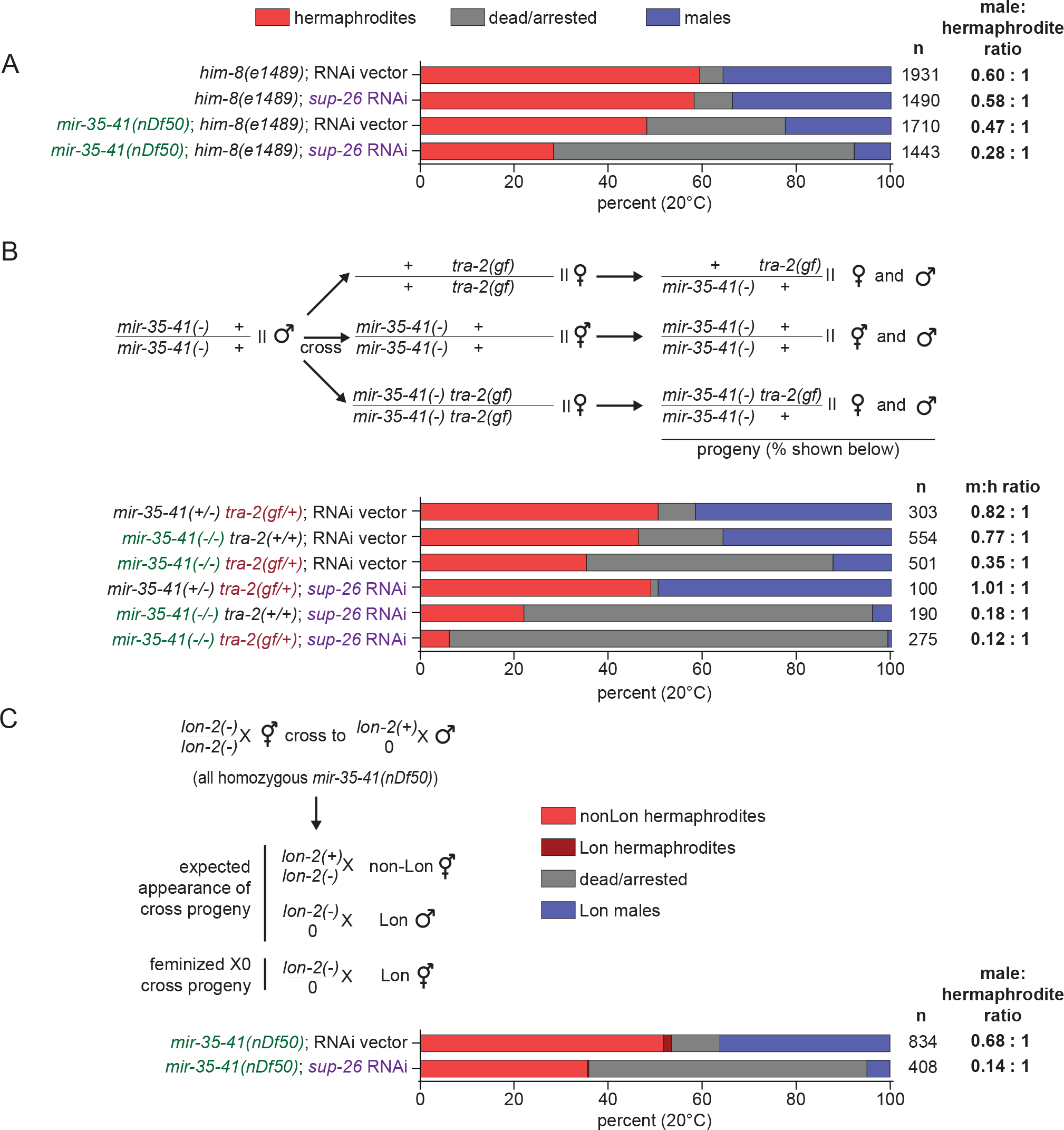
Feminizing mutations preferentially enhance *mir-35-41(nDf50)* lethality in males. A) Percent dead/arrested, male or hermaphrodite progeny in a high-incidence-of-males (*him*) background. *Sup-26(RNAi)* reduces the proportion of males among surviving animals. B) Top: Schematic of crosses performed to generate progeny scored in graph. The crosses were conducted on RNAi plates, so RNAi affects the maternal and zygotic contribution of *sup-26* in the progeny. Males of the same genotype were used for all crosses in order to compare equal dosage of *tra-2(e2020gf)* across different *mir-35-41* genotypes. Colored text highlights functional genetic differences between genotypes. *Sup-26(RNAi)* and *tra-2(e2020gf)* preferentially enhance male lethality in *mir-35-41(nDf50)*. C) Top: Schematic of cross with recessive X-linked marker (*lon-2(e678)*) to assess potential somatic feminization of XO cross progeny. Bottom: Percent of progeny. The rare Lon hermaphrodites likely represent self-progeny (also see methods). In addition to progeny shown, two males were scored as non-Lon on empty vector RNAi. *Sup-26(RNAi)* does not increase the apparent proportion of somatically feminized XO animals (Lon hermaphrodites).

Because *sup-26(lf)* and *tra-2(e2020gf)* are feminizing mutations, the reduced number of males observed in *mir-35-41(nDf50)*; *him-8(e1489)* on *sup-26(RNAi)* and in *tra-2(e2020gf)* could be a result of feminization of males rather than lethality. In this case, feminized XO animals would be spuriously counted as hermaphrodites based on morphology. (Additionally, *mir-35-41(nDf50)*; *him-8(e1489)* males are abnormal, indicating that *mir-35-41(nDf50)* may be a genetic background that is sensitized to feminization (McJunkin and Ambros 2014).) To distinguish between feminization and preferential male lethality, *mir-35-41(nDf50)* hermaphrodites containing a recessive X-linked marker (*lon-2(e678)*) were crossed to non-Lon *mir-35-41(nDf50)* males. If *sup-26(RNAi)* causes feminization of *mir-35-41(nDf50)*; *him-8(e1489)* males, then an increase in Lon hermaphrodite-like progeny would be observed. *Sup-26(RNAi)* did not increase the proportion of Lon hermaphrodites, indicating that the reduced number of males observed is due to an increase in embryonic/early larval lethality among males and not feminization of XO animals (Figure 3C).

### Male preferential lethality is linked to karyotype

The preferential lethality of males in feminized *mir-35-41(nDf50)* animals examined thus far could be a result of the males’ XO karyotype or due to the male developmental program downstream of the sex determination pathway. To distinguish between these two possibilities, we decoupled XO karyotype from male development using a *fem-3(e1996null)* mutation. In this background, both XX and XO animals develop as females. To distinguish between morphologically similar XO and XX animals, we employed a crossing strategy in which XX progeny were marked by an mCherry transgene integrated on the X chromosome (Figure 4A). If lethality in a feminized *mir-35-41(nDf50)* background still preferentially affects XO animals even when they develop as females, then this preferential lethality is due to XO karyotype and not because of male development.

**Figure 4.**
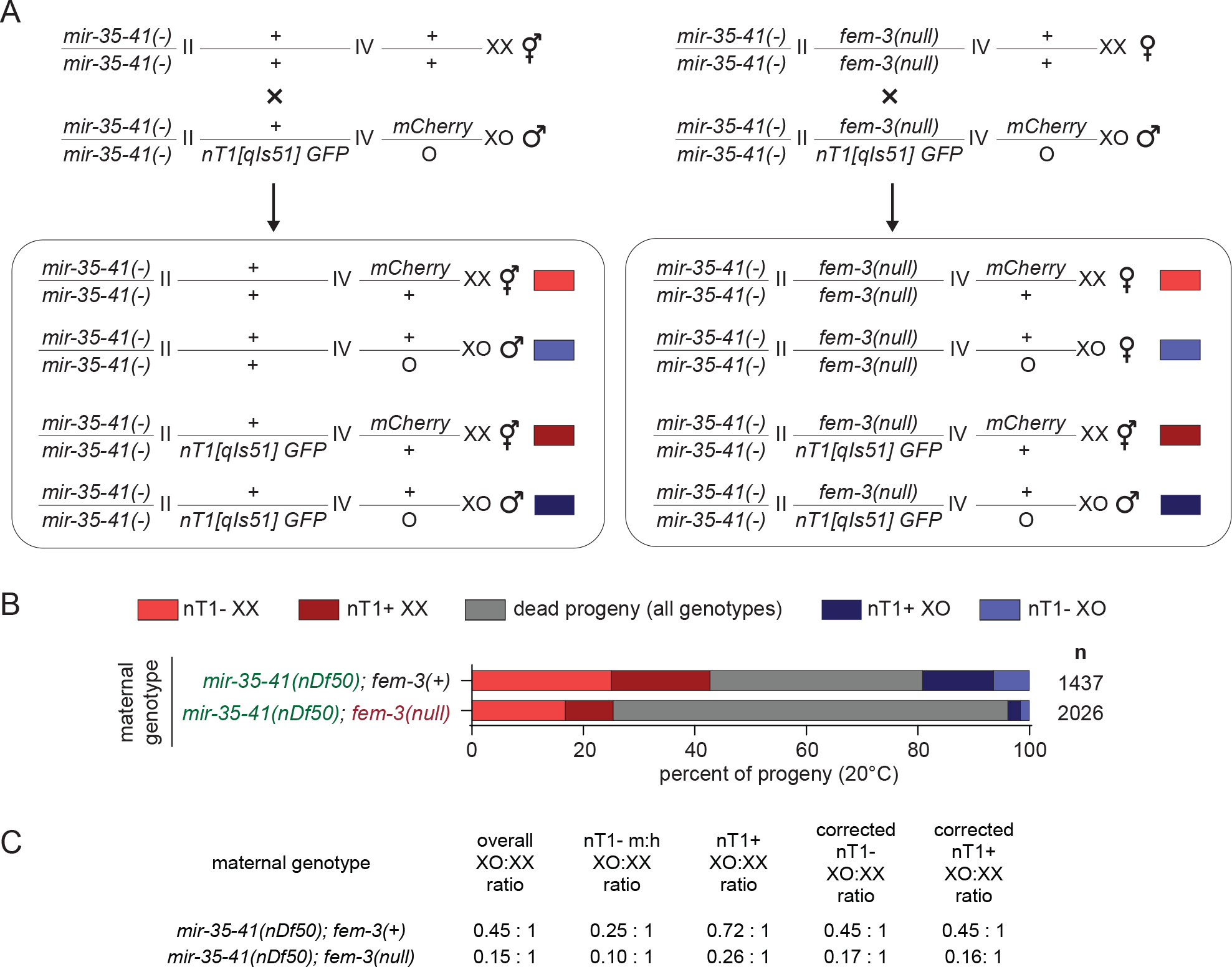
XO karyotype underlies preferential death of males in feminized *mir-35-41(nDf50)* animals. A) Schematic of cross. Males contain a GFP-marked nT1 balancer and an X-linked mCherry transgene which aid in distinguishing *fem-3* genotype and number of X chromosomes in progeny. B) Percent of progeny from crosses in each category. Dead/arrested embryos and larvae were not scored for fluorescent markers, and thus are likely a mixture of genotypes. C) XO:XX ratios in populations of progeny from crosses. Corrected values assume that differences in XO:XX ratio in nT1+ and nT1− population in the control *mir-35-41(nDf50); fem-3(wild type)* cross are due to non-Mendelian segregation of the nT1 balancer.

Overall, progeny resulting from the cross involving *mir-35-41(nDf50)*; *fem-3(null)* mothers showed greater lethality than progeny from the *mir-35-41(nDf50)* mothers (Figure 4B). The genotypes of the dead embryos are not defined since embryos may die before expressing fluorescent markers. When examining the live *mir-35-41(nDf50)*; *fem-3(null)* progeny (those that lack the nT1 balancer chromosome), the XO:XX ratio was lower than that in the comparable nT1-*mir-35-41(nDf50)* progeny (0.10 compared to 0.25) (Figure 4B-C). This indicates that, as in previous experiments, the feminizing mutation (*fem-3(null)*) preferentially enhanced lethality among XO animals. However, here, the *mir-35-41(nDf50); fem-3(null)* XO animals develop as females, not males. Therefore, the preferential enhancement of lethality is due to the XO karyotype since it is manifested in the absence of a male developmental outcome.

While the overall ratio of males to hermaphrodites was 0.45:1 in the control *mir-35-41(nDf50)* cross, this ratio was lower among animals lacking the nT1 balancer (0.25) and higher among those containing the nT1 balancer (0.72) (Figure 4C). Though this may seem like a surprising discrepancy due to the fact that nT1 is not balancing any mutant alleles in this cross, the disparity can be explained by the previously-observed tendency for nT1 to segregate preferentially with nullo-X sperm (Edgley *et al.* 2006). If we assume that this segregation preference accounts for the difference between the XO:XX ratio in nT1+ animals and nT1-animals in the control cross, then we can determine a correction factor for this preference (a coefficient for each value that adjusts it to the average XO:XX ratio for the cross). We can then apply the same coefficients to the XO:XX ratios of the *mir-35-41(nDf50)*; *fem-3(null)* cross to correct for biases resulting from nT1 segregation alone (see below).

When examining nT1+ progeny of these crosses, *mir-35-41(nDf50)*; *fem-3(null)/*nT1 progeny show a more skewed sex ratio (0.26) compared to *mir-35-41(nDf50);* +/nT1 progeny (0.72). In fact, after correcting the XO:XX ratios for skewed segregation of nT1, the preferential death of males occurs at the same rate in nT1+ progeny of this cross as among nT1-progeny (both 0.16-0.17). The most plausible interpretation of this finding is that the maternal *fem-3* genotype has a greater impact than the zygotic *fem-3* genotype on this phenotype since both nT1+ and nT1− progeny lack maternal *fem-3* (while they differ in their zygotic dose of *fem-3*). Thus, observing the same effect size in nT1+ and nT1− progeny suggests that the enhancement of XO lethality by *fem-3* loss of function occurs via a maternal effect.

### Maternal germline feminization preferentially enhances lethality in males

All our results thus far indicate that when *mir-35-41(nDf50)* is combined with feminizing genetic backgrounds (*sup-26(RNAi)*, *tra-2(e2020gf)*, *her-1(null)*, *fem-3(null)*), higher rates of lethality occur, and that this enhanced lethality preferentially affects XO animals. Because *sup-26(RNAi)*, *her-1(null)*, *fem-3(null)* all affect both the soma and the germline (Ellis and Schedl 2007; Wolff and Zarkower 2008; McJunkin and Ambros 2017), and *tra-2(e2020gf)* has a much greater effect on the germline than the soma (Doniach 1986), we sought to distinguish whether these effects are mediated by the somatic or the germline sex determination pathway. To this end, we tested feminizing mutations in genes that act only in the germline sex determination pathway for their effect on *mir-35-41(nDf50)* lethality. First, a germline feminizing lesion, *fog-2(null)*, was introduced to the *mir-35-41(nDf50)* background. Simple male-female crosses demonstrated that *fog-2* loss of function enhanced preferential death of males in *mir-35-41(nDf50)*, indicating that the germline-specific sex determination pathway is implicated in this phenotype (Figure 5A).

**Figure 5.**
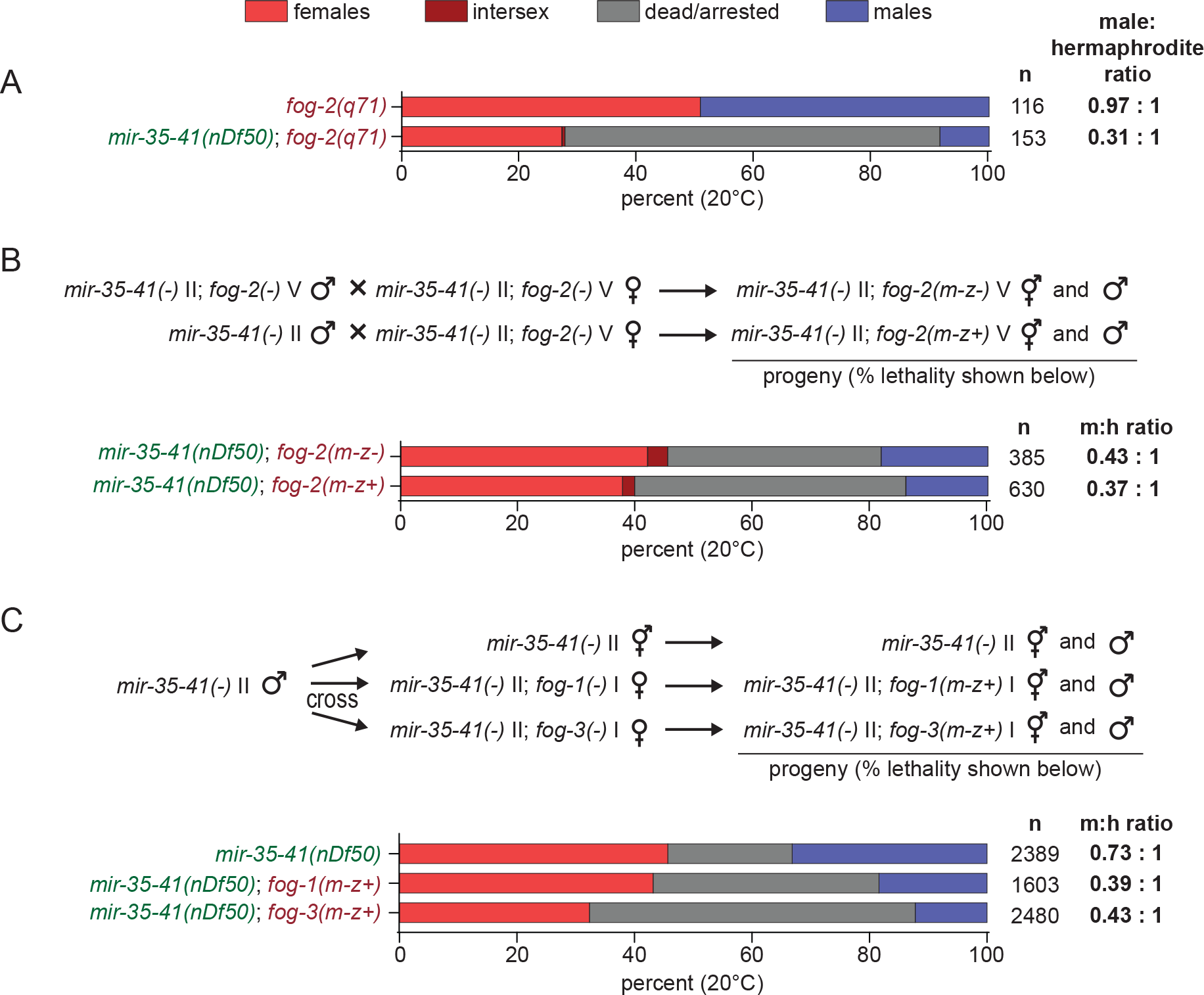
Feminization of the germline causes preferential death of *mir-35-41(nDf50)* males, and this is a maternal effect. A) Percent dead/arrested, male or female progeny in a *fog-2(q71)* background, with wild type or deleted *mir-35-41*. B) Percent dead/arrested, male or female progeny. Top bar: both parents are *mir-35-41(nDf50)*; *fog-2(q71)*. Bottom bar: mother is *mir-35-41(nDf50)*; *fog-2(q71)*. Father is *mir-35-41(nDf50); fog-2(wild type)*. C) Top: Schematic of cross. Males also contained a GFP integrated transgene to prevent the scoring of self progeny. Bottom: Percent dead/arrested, male or female progeny.

We next sought to determine whether the effect of the germline-specific sex determination pathway on this phenotype is via maternal effect. Therefore, we crossed *mir-35-41(nDf50)* males to *mir-35-41(nDf50)*; *fog-2(null)* females and scored viability of the progeny. Like animals lacking both maternal and zygotic *fog-*2, mir*-35-41(nDf50)*; *fog-2(m-z+)* animals showed preferential death among males, suggesting that the perturbation of germline sex determination pathway enhances this phenotype via a maternal effect (Figure 5B). These animals have normal zygotic germline sex determination. Thus, maternal germline feminization likely underlies the preferential male lethality phenotype. This is consistent with the apparent maternal effect of *fem-3(null)* on enhanced lethality and preferential male lethality in *mir-35-41(nDf50)* and the previously observed partial maternal effect of the *mir-35-41* family on sex determination and other phenotypes (Alvarez-Saavedra and Horvitz 2010; McJunkin and Ambros 2014, 2017).

To determine whether other players in the canonical germline-specific sex determination pathway were also implicated in this phenotype, we tested the effect of maternal loss of *fog-1* or *fog-3* function in the *mir-35-41(nDf50)* background. Like *fog-2*, loss of maternal *fog-1* or *fog-3* enhanced lethality and male-preferential lethality of *mir-35-41(nDf50)*, with *fog-3(null)* having a stronger effect than *fog-1(null)* (Figure 5C). Thus, the canonical germline sex determination pathway is involved in preventing the enhanced lethal phenotypes, and *mir-35-41(nDf50)* mutants are sensitized to maternal germline feminization, especially among XO males.

## Discussion

We found that the *mir-35* family, which regulates sex determination in the soma and germline, also has unexpected lethal interactions with the germline sex determination pathway. In the *mir-35-41(nDf50)* background, multiple germline feminizing mutations enhance lethality of both sexes, while preferentially enhancing that of XO males. To our knowledge, this is the first synthetic lethal effect of the germline sex determination pathway. In some cases, assigning significance to the enhancement of lethality in a genetic background that alone elicits lethality can be difficult. However, two factors: (1) the consistency of enhancement by feminizing mutations and failure to enhance by masculinizing mutations and (2) the preferential effect on XO males both indicate that we are observing a specific synthetic phenotype and genetic interaction of two pathways.

What is the basis of this synthetic lethality? One possibility is that *mir-35-41(nDf50)* mutants with feminizing mutations experience conflicting signals through the sex determination pathway due to the masculinization caused by derepression of multiple target genes downstream of *mir-35-41*. Such conflicting developmental signals could be deleterious. However, this is counterintuitive in light of the fact that the sex determination pathway is fairly linear, and legacy mutations show clear epistatic effects, rather than additive parallel genetic interactions. However, genetic data shows that *mir-35-41* exerts a partially maternal effect on sex determination (McJunkin and Ambros 2017). Our working model is that *mir-35-41* acts as a developmental timer, preventing premature sex-specific gene expression. According to this model, removal of *mir-35-41* function could disrupt the clean epistatic and linear genetic relationships of the sex determination pathway, since the order of events in the pathway would be disrupted. Thus, the incoherence of premature male gene expression in *mir-35-41* could conflict with feminizing mutations.

Another model is that feminizing mutations in the sex determination pathway ameliorate aberrant masculinization of *mir-35-41(nDf50)* mutants, while having a synthetic lethal effect with another pathway misregulated in *mir-35-41(nDf50)*. This is consistent with the concept that the *mir-35-41(nDf50)* family controls multiple targets, pathways, and phenotypes (Alvarez-Saavedra and Horvitz 2010; Liu *et al.* 2011; Massirer *et al.* 2012; McJunkin and Ambros 2014, 2017; Kagias and Pocock 2015). This latter model would imply that the germline sex determination pathway controls other aspects of development outside of sex determination. FOG-1 and FOG-3 bind to the mRNA of 81 and ~1000 target genes, respectively (Noble *et al.* 2016; Aoki *et al.* 2018). While regulation of many of these targets must mediate the sex determination function of FOG-1 and FOG-3, regulation of some targets may also contribute to other biological pathways we have yet to understand. Mutation of a fog gene may require homeostatic changes in downstream non-sex determination genes to result in a clean germline feminization phenotype without deleterious pleiotropic effects. The *mir-35* family may be important in buffering these homeostatic changes, thus revealing potentially deleterious consequences in the microRNA family mutant.

The above models provide frameworks for conceptualizing the generalized enhanced lethality caused by feminizing mutations in *mir-35-41(nDf50)*, but what could be the basis of the preferential effect on XO animals? Because the enhanced lethality preferentially affects animals with an XO karyotype more than those with an XX karyotype, they may result from perturbation of chromatin modifying complexes that bind and modulate gene expression on the X chromosome. One of these complexes is the DCC. In wild type animals, the DCC is stable and loaded onto X chromosomes only in XX animals (Meyer 2005). Aberrant activation of the DCC in XO animals causes XO-specific lethality. A feedback loop whereby TRA-1 reinforces repression of *xol-1* transcription could explain some of the genetic effects seen here, though not those of *fog-1* and *fog-3* (Hargitai *et al.* 2009). While an effect on the DCC is one possible model for the phenotype we observe here, two observations oppose this model. First, the *mir-35-41(nDf50)* mutation causes masculinization of gene expression, which would be expected if the DCC were inactivated, not aberrantly activated. Second, the enhancement of lethality by feminizing mutations observed here importantly affects both XX and XO (with a stronger effect on XO) animals; this would be somewhat unusual for a DCC-activating mutation since DCC activation in XX animals is tolerated and essential.

A second system that modifies X-linked chromatin differently than autosomes is the maternal effect sterile (MES) proteins. MES-2/3/6 make up the polycomb repressive complex 2 (PRC2) in *C. elegans* and are critical in silencing the X chromosome in the germline via H3K27 methylation (Bender *et al.* 2004). Their activity is opposed by MES-4, which modifies autosomes with activating H3K36me and excludes autosomal binding by MES-2/3/6 (Bender *et al.* 2006). Many of the synthetic multivulva class B (synMuv B) genes are required to contain the germline activity of the MES proteins to their proper compartment. In certain synMuv B mutants, including inactivation of *lin-35*/Rb, somatic cells take on a germline-like gene expression signature which leads to early larval arrest at high temperature (Petrella *et al.* 2011; Wu *et al.* 2012). In these contexts, loss of function of either MES-2/3/6 or MES-4 suppresses larval arrest and germline-like gene expression in the soma. Whether this larval arrest could preferentially affect XO animals is not known, but this would be a reasonable prediction since the arrest is due to a germline-like chromatin environment in the soma (and thus somatic X chromosome silencing).

In *mir-35-41(nDf50)* mutant embryos, *lin-35* does not accumulate to wild type levels, leading to a *lin-35(lf)*-like enhanced RNAi phenotype (Massirer *et al.* 2012). Since *lin-35* also promotes female development, the low *lin-35* level in *mir-35-41(nDf50)* could also underlie the observed masculine gene expression pattern (Grote and Conradt 2006). If low *lin-35* in *mir-35-41(nDf50)* also causes a germline-like chromatin state in the soma, this could possibly account for the lethality phenotype observed here, including the deleterious effect on both sexes as well as the XO-preferential effect.

Future studies should attempt to distinguish between these models. Is the male-like gene expression of *mir-35-41(nDf50)* required for the enhanced lethality and preferential death among XO? Is a germline-like chromatin state present in *mir-35-41(nDf50)*? Is the low level of *lin-35* responsible for these phenotypes? Delineating the pathways responsible for these phenotypes downstream of *mir-35-41* will be facilitated by CRISPR/Cas9-mediated genome editing. By relieving single target genes from *mir-35-41*-mediated repression, we can begin to understand which phenotypes are downstream of each axis of the pathway, and thus eventually sort out which gene expression changes and phenotypes are causative of each other or genetically separable.

## Acknowledgements

Thank you to Victor Ambros in whose lab this work was initiated. Thank you to Rosalind Lee for strains and ongoing support. We acknowledge Eric Haag for critical reading of the manuscript. The McJunkin lab is funded by the NIDDK Intramural Research Program (1ZII DK075147-01). Some strains were provided by the CGC, which is funded by NIH Office of Research Infrastructure Programs (P40 OD010440).

---

**Figure S1.**
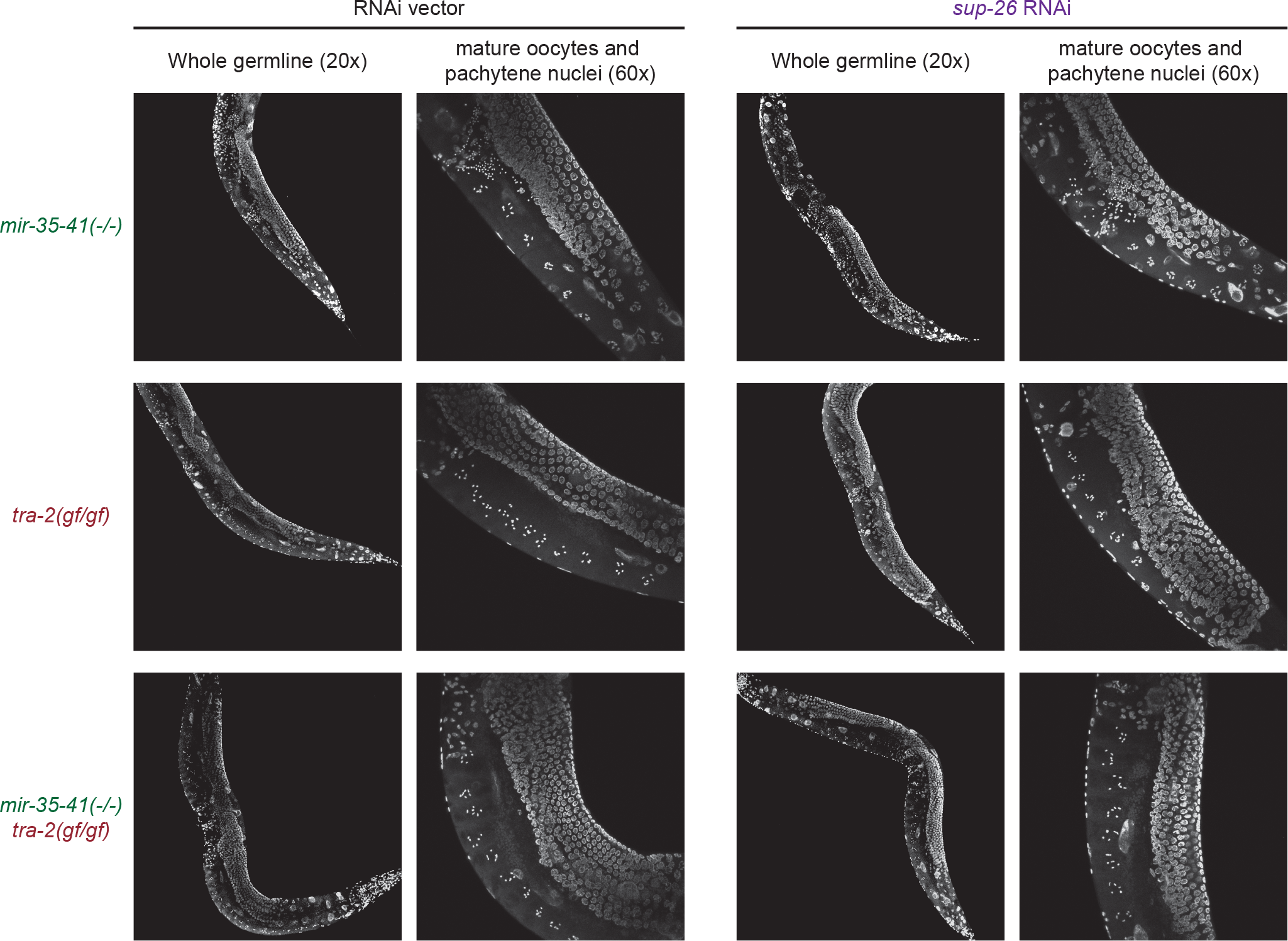
Germline development is grossly normal in *mir-35-41(nDf50)*, *tra-2(e2020gf)*, and *mir-35-41(nDf50) tra-2(e2020gf)* hermaphrodites or females. DAPI staining of first-day-gravid adults of indicated genotypes. RNAi treatment and genotypes are identical to mothers whose progeny are scored in Figure 2A. All genotypes show normal oocyte development.

**Figure S2.**
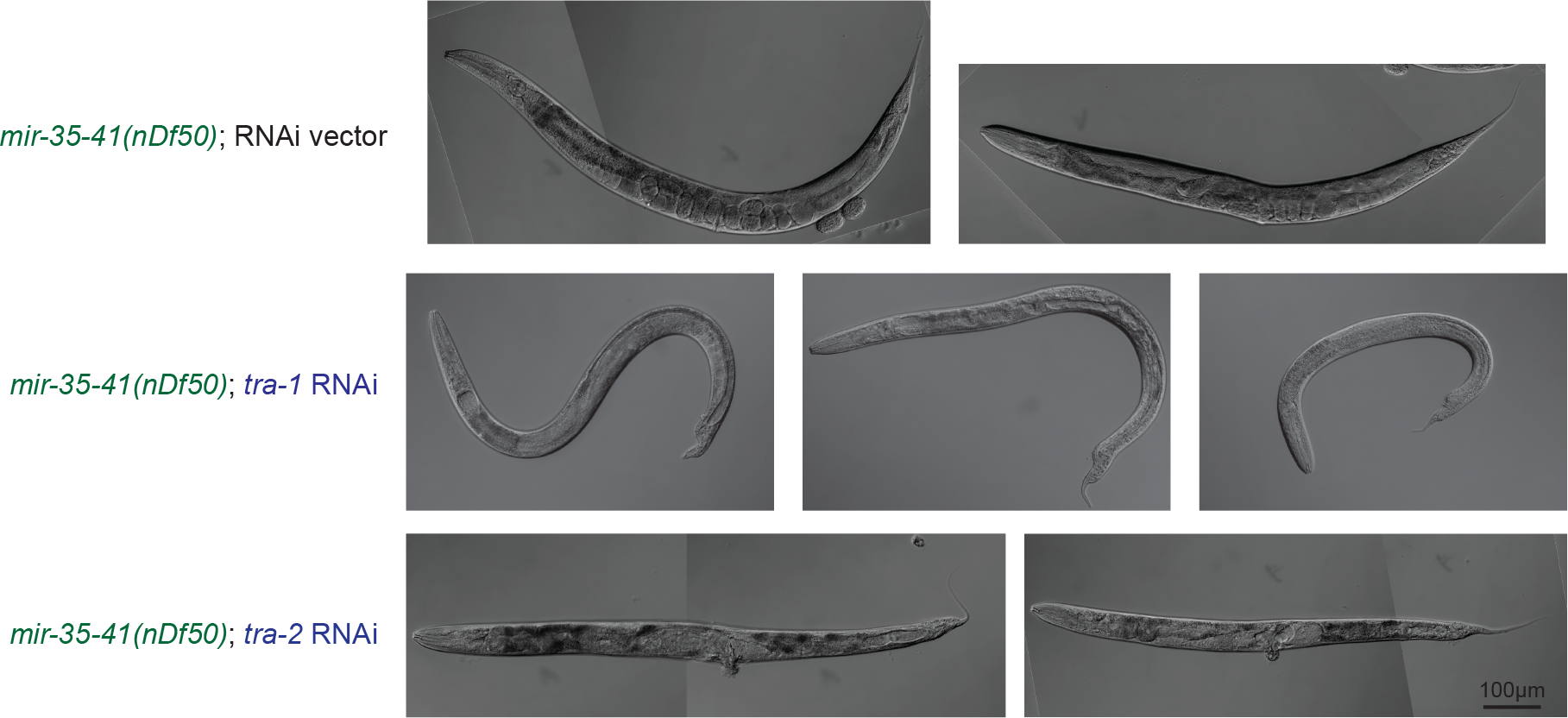
RNAi against tra-1 and tra-2 elicits expected transformed and intersex phenotypes. DIC micrographs of surviving animals quantified in Figure 2C.

